# The human gut virome database

**DOI:** 10.1101/655910

**Authors:** Ann C. Gregory, Olivier Zablocki, Allison Howell, Benjamin Bolduc, Matthew B. Sullivan

**Author notes:** Correspondence to: Matthew Sullivan. These authors contributed equally (Ann C. Gregory, Olivier Zablocki).

## Abstract

The gut microbiome profoundly impacts human health and disease, but viruses that infect these microbes are likely also important. Problematically, viral sequences are often missed due to insufficient reference viral genomes. Here we (i) built a human gut virome database, GVD, from 648 viral particle metagenomes or microbial metagenomes from 572 individuals previously searched for viruses, (ii) assessed its effectiveness, and (iii) conducted meta-analyses. GVD contains 13,203 unique viral populations (approximately species-level taxa) organized into 702 novel genera, which roughly doubles known phage genera and improves viral detection rates over NCBI viral RefSeq nearly 60-fold. Applying GVD, we assessed and rejected the idea of a ‘core’ gut virome in healthy individuals, and found through meta-analyses that technical artifacts are more impactful than any ‘treatment’ effect across the entire meta-study dataset. Together, this foundational resource and these findings will help human microbiome researchers better identify viral roles in health and disease.

## Main text

The human gut microbiome is now thought to play an integral role in health and disease^1–4^. Persistent alterations in the structure, diversity and function of gut microbial communities—dysbiosis— are increasingly recognized as key contributors in the establishment and maintenance of a growing number of disease states^5–7^, including obesity^8^ and cancer^9^. Gut dysbiosis can develop from complex interplays between host, cognate microbiota and external environmental factors^10,11^. Within the gut microbial consortium, the bacteriome has been the most extensively studied, where significant shifts in population dynamics have been observed between healthy and diseased individuals^12^. However, emerging views^10,13,14^ suggest that the gut virome plays an important role in homeostatic regulation and disease progression through multiple interaction paths with the co-occurring bacteriome, and even directly with human immune system components^15^.

The first step in studying viruses in complex communities is to “see” them. Problematically, identifying viral sequences in large datasets is notoriously challenging. Because viruses lack a universal viral marker^16^, as opposed to bacterial 16S rRNA for example, researchers often resort to sequence homology searches against reference databases (e.g. NCBI viral RefSeq). Such searches are variably successful with anywhere from 14% to 87% of the observed gut viral genomes having detectable similarity to viruses in such databases^10^. This large range stems from several factors that are not mutually-exclusive including the following: (i) broad under-representation of viral genome space in databases, (ii) non-standardized database usage per study, (iii) overrepresentation of certain virus groups due to sample preparation and cultured host availability, and (iv) natural sample variation. In addition, although viral reference datasets are being generated at unprecedented rates^17^, these new data are rarely incorporated for cross-comparisons, which inflates virus novelty in new datasets and/or leaves many virus sequences undetected. Therefore, given the rapid accrual of so many studies, there is a need to aggregate their findings into a central gut-specific database to improve gut virome inference capabilities.

Here we collected and curated 648 gut metagenomes from 21 datasets (i.e., any metagenomic dataset that looked at gut viruses published before 2018), consistently processed them to map known and unknown viral populations, and used this in multiple meta-analyses to assess improvement and reveal new biology. The resulting Gut Virome Database (GVD) was born by (i) collecting 648 gut metagenomes from 572 individuals, (ii) extensive metadata curation through literature mining and, as needed, direct communication with the original researchers, and (iii) re-analysis of the virome data to establish consistent processing and extensive virus identification. The value of GVD was assessed for performance against the best currently available databases (NCBI viral RefSeq and IMG/VR^18^), and then used to re-evaluate global diversity patterns and the relationship between gut virome diversity and diet.

## RESULTS AND DISCUSSION

### GVD contains 13,204 viral populations, dominated by phages

To build a collection of the commensal human gut virome, 648 metagenomic samples from 572 individuals were processed from all datasets publicly available as of December 2017 (n=19), along with 2 unpublished datasets where access was granted prior to publication. These studies represented a total of 1.28 Tbp of sequence data derived from a spectrum of gut virome study areas including: (i) healthy gut viromes of infants^19,20^ and adults^21–26^, as well as individuals experiencing (ii) fecal matter transplant, or FMT^27–31^, (iii) inflammatory bowel disease, or IBD^32,33^, (iv) HIV infection^34^, (v) Type I diabetes^35,36^, (vi) malnutrition^37^, or (vii) chronic fatigue syndrome^38^ (see **Supplementary Table 1**). Datasets had a worldwide distribution, though most originated from the United States (48.4%; **Fig. 1a**). All reads were processed consistently, assembled into contigs and viral-like sequence were identified using three independent methods and validated by cross-comparisons between methods (**Fig. 1b**, see Methods). To avoid duplicate viral fragments/partial virus genomes across the datasets, contigs were de-replicated by clustering sequences according to percentage of average nucleotide identity (ANI) and sequence length. Multiple reports^17,39–43^ have revealed that > 95% ANI was a suitable threshold for defining a set of closely-related discrete ‘viral populations’, with follow-on studies suggesting that this cut-off establishes populations that are largely concordant with a biologically relevant viral species definition^39,41,44^. Using this clustering strategy, we identified highly variable numbers of unique viral populations per study (range: 0 - 3596; mean = 670) (**Supplementary Fig. 1a**). GVD comprises 13,203 viral populations (N50 = 34,220 bp; L50 = 2,066 bp). For context, NCBI’s viral RefSeq v88 (released May 2018) database holds 8,013 viruses of eukaryotes, bacteria and archaea from all environments, combined. Moreover, if only comparing phage genomes to the same database, GVD contains 7 times more phages compared to the entire set of cultured phage isolates in viral RefSeq to date. Thus, GVD greatly augments the repertoire of known viruses in the human gut.

**Figure 1.**
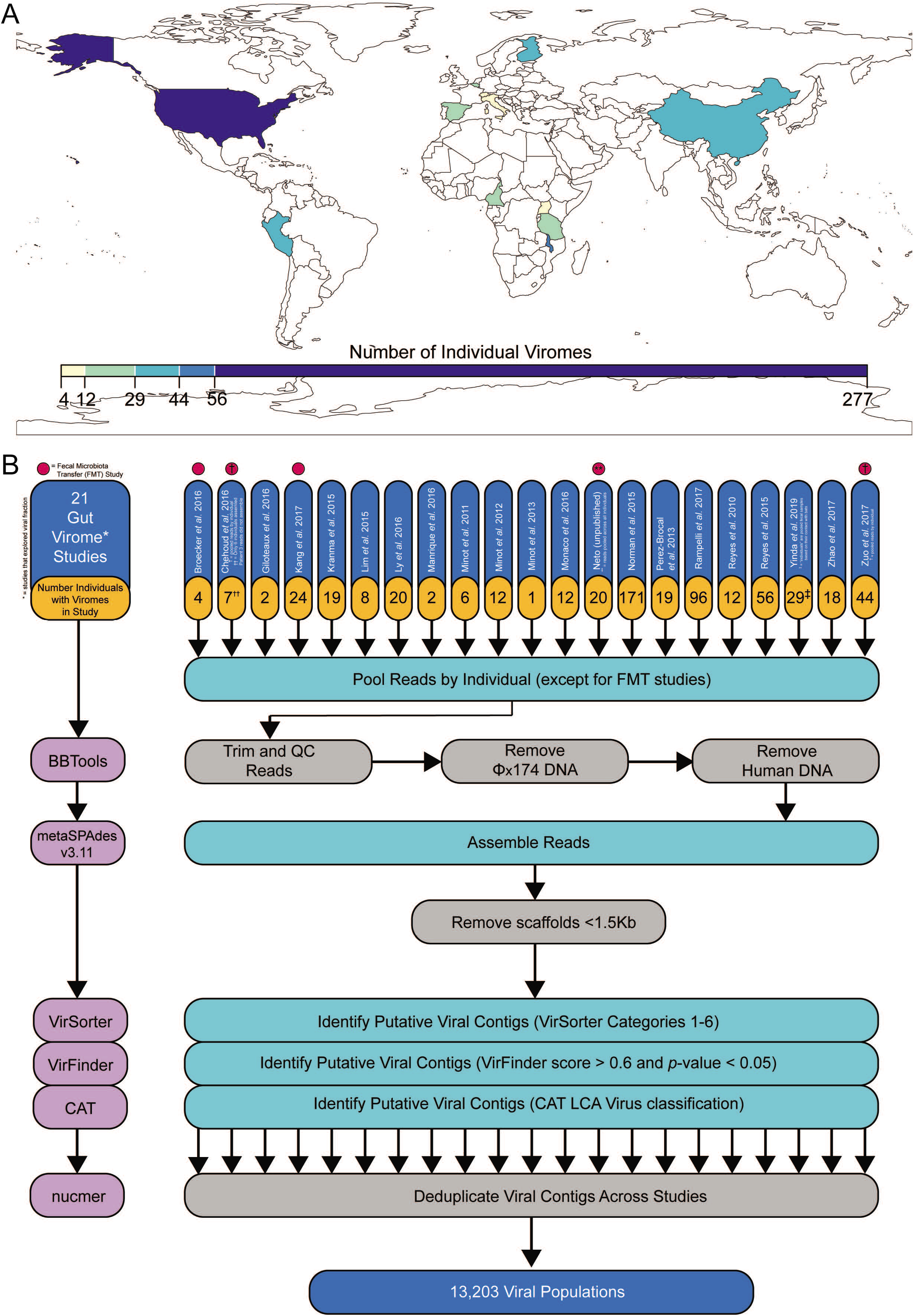
Overview of studies and meta-analyses comprising the Gut Viral Database (GVD). **(a)** Global heatmap of the world showing the number of individual’s gut viromes coming from different countries within the GVD. Importantly, individual’s viromes coming from the Cameroon were pooled based on their location, age, and contact with bats. The pools were counted as a single individual’s virome for our analyses. **(b)** Pipeline for the selection and processing of human gut virome datasets (see **Methods**). Datasets were processed individually and, within each dataset, viromes were pooled by individual, except for fecal microbiota transfer (FMT) studies and data that was given to us prior to publication (Yinda et al., 2019; Neto et al. (unpublished)). Reads were filtered for quality and trimmed and reads that mapped to Φx174 and the human genome were removed. The remaining reads were assembled into scaffolds, filtered for lengths ≥1.5kb, and run through tools that collectively utilize homology to viral reference databases, probabilistic models on viral genomic features, and viral *k*-mer signatures to identify viral contigs. Viral contigs were then deduplicated to get a total of 13,203 viral populations.

Taxonomically, 96.1% of GVD viral populations are bacterial viruses (i.e., phages), with a minority of GVD viral populations more likely to represent eukaryotic viruses (3.8%) and archaeal viruses (0.1%) (**Fig. 2a**). Though in the minority, the 505 eukaryotic viruses were taxonomically diverse (14 families), dominated by ssDNA families *Anelloviridae* (72%), *Genomoviridae* (10%) and *Circoviridae* (8%). All, with the exception of Genomoviruses, have been reported previously in the datasets underlying GVD^34^. Among the phages, 82% did not have ICTV classification, with the remaining fraction comprised of dsDNA tailed phage families (*Siphoviridae, Myoviridae* and *Podoviridae*), *Microviridae* and *Inoviridae* (see **Supplementary Table 2**). Twelve unknown archaeal viral populations were detected, with no close genome/gene homology to any of the classified archaeal viruses. The high number of unclassified phages likely results from underrepresentation of gut phages in the database, coupled to unresolved and/or missing taxonomic assignments for ~ 60% of reference phage genomes in RefSeq, with the currently classified fraction organized into ~250 genera^45^. To fill this phage and archaeal virus taxonomic classification gap, we used a genome-based, gene-sharing network strategy^46,47^ that *de novo* predicts genus-level groupings (‘viral clusters’ or ‘VCs’) from viral population data. A network was computed from 6,373 GVD phage genomes (only those ≥10 kb in length; 48% of GVD), combined with 2,304 curated reference phage genomes from NCBI Viral RefSeq (version 88). The resulting gene-sharing network (**Fig. 2b**) revealed 957 VCs, 702 of which were novel and exclusively composed of GVD genomes (3,220 viral genomes or ~51% of GVD genomes). This would roughly double the current number of ICTV-recognized phage genera. Though not explored here, as our goals focused on taxonomic classification, the shared protein content within and between VCs calculated in our network analyses could be used to guide qPCR assays for NGS validation^48^ and/or tracking of viruses at either the viral population- or genera-level under changing conditions^35^.

**Figure 2.**
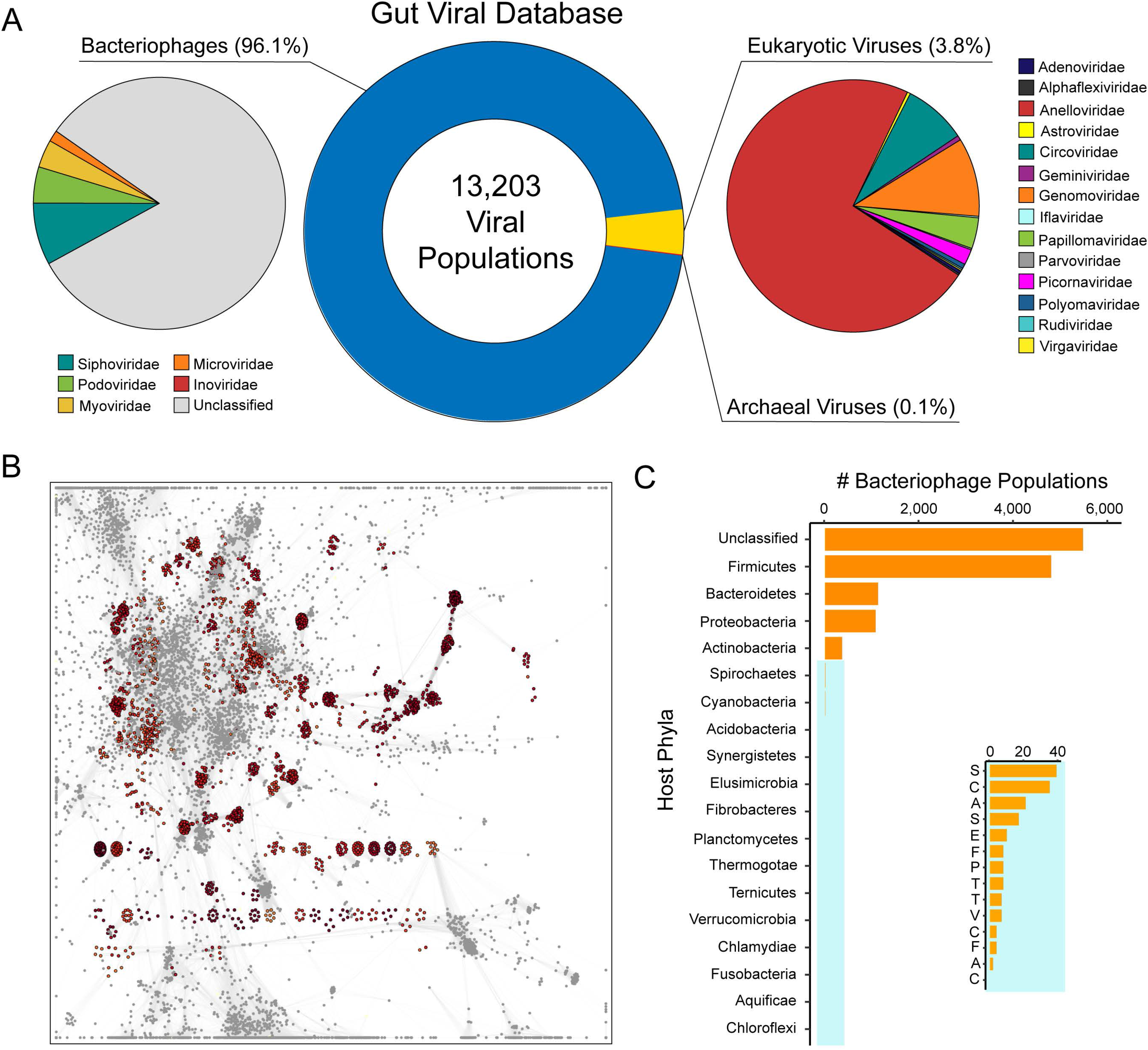
The Gut Viral Database (GVD). **(a)** Pie charts showing the number of bacteriophages, eukaryotic viruses, and archaeal viruses in the GVD (center) and their familial taxonomic composition by the bacteriophages (left) and the eukaryotic viruses (right). **(b)** Gene-sharing taxonomic network of the GVD, including viral RefSeq viruses v88. RefSeq viruses are highlighted in red. Every node represent a virus genome, while connecting edges identify significant gene-sharing between genomes, which form the basis for their clustering in genus-level taxonomy. **(c)** Bar chart showing the number of bacterial host phyla of the GVD bacteriophages, with an inset providing resolution for the low frequency bacteria host phyla. Putative host phyla per each bacteriophage population are in **Supplementary Table 3**

Next, we sought to link phage populations to their hosts using *in silico* strategies (see **Methods**). The most common identifiable phage hosts (**Fig. 2c**) in GVD belonged the bacterial phylum Firmicutes (38%), about 2-fold more than the next most abundantly identified host phyla (Bacteroidetes and Proteobacteria; see **Supplementary Table 2**). Though Firmicutes and Bacteroides are the most prominent bacterial phyla in the human gastrointestinal tract^49^, Firmicutes typically outnumber Bacteroidetes in unhealthy individuals with metabolic and digestive disorders^50–52^. GVD metagenomes originated from ~16% healthy individuals and ~84% unhealthy individuals, many of which have metabolic and digestive disorders. Thus, it is perhaps not surprising that most of the annotated viral populations were linked to the phylum Firmicutes.

### GVD significantly improves virus detection in all gut datasets

We then quantitatively evaluated virus identification sensitivity (through read mapping) between multiple databases by comparing the number of identified viral populations in each study detected by GVD, viral RefSeq v88, IMG/VR 1.1 (2018 release) and the individual virome datasets (‘IV’) from each study (**Fig. 3**). For the latter, IV reads were mapped against viral populations (predicted in this study) derived exclusively from its matching IV. In all datasets, GVD surpassed viral RefSeq (mean increase: 59-fold ± 95-fold) and IVs (mean increase: 3.2-fold ± 6.6-fold). In 5 of 18 studies (28%), GVD outperformed IMG/VR (mean increase: 1.1-fold ± 2-fold), with the remaining studies finding no significant difference between or too low of a sample size to compare GVD and IMG/VR. After GVD, IMG/VR was the next best performing database for viral detection in the gut, as our tests showed an average of 49-fold (± 87-fold) increase over viral RefSeq. IMG/VR was expected to surpass viral RefSeq, as it aggregates both cultivated reference virus genomes, >12,000 prophages and >700,000 uncultivated virus genomes/fragments from many environments, including multiple human body sites^53^. Moreover, given the high performance of IMG/VR in our tests, we wondered about the extent of viral population overlap with GVD (**Fig. 3b**). There were 1,730 viral populations shared between the two databases, but still each database is overwhelmingly unique (82% and 69% unique to GVD and IMG/VR, respectively). This is because IMG/VR includes human gut studies that did not explore the viral fraction as well. Overall, the significant increase in virus detection by GVD over other databases (two-tailed Mann-Whitney U-tests; *p*-value < 0.05) highlights the low representation of gut viruses recorded in RefSeq and thus demonstrates the value of GVD for sequence-based virus identification in human gut microbiome datasets. Because the datasets used to compile GVD were originally analyzed most often (55% of the studies) using viral RefSeq as the primary source to identify viruses (**Supplementary Table 1**), we wondered whether significant fractions of viruses could have been missed, and whether a possibly reduced viral “signal” would influence previous conclusions.

**Figure 3.**
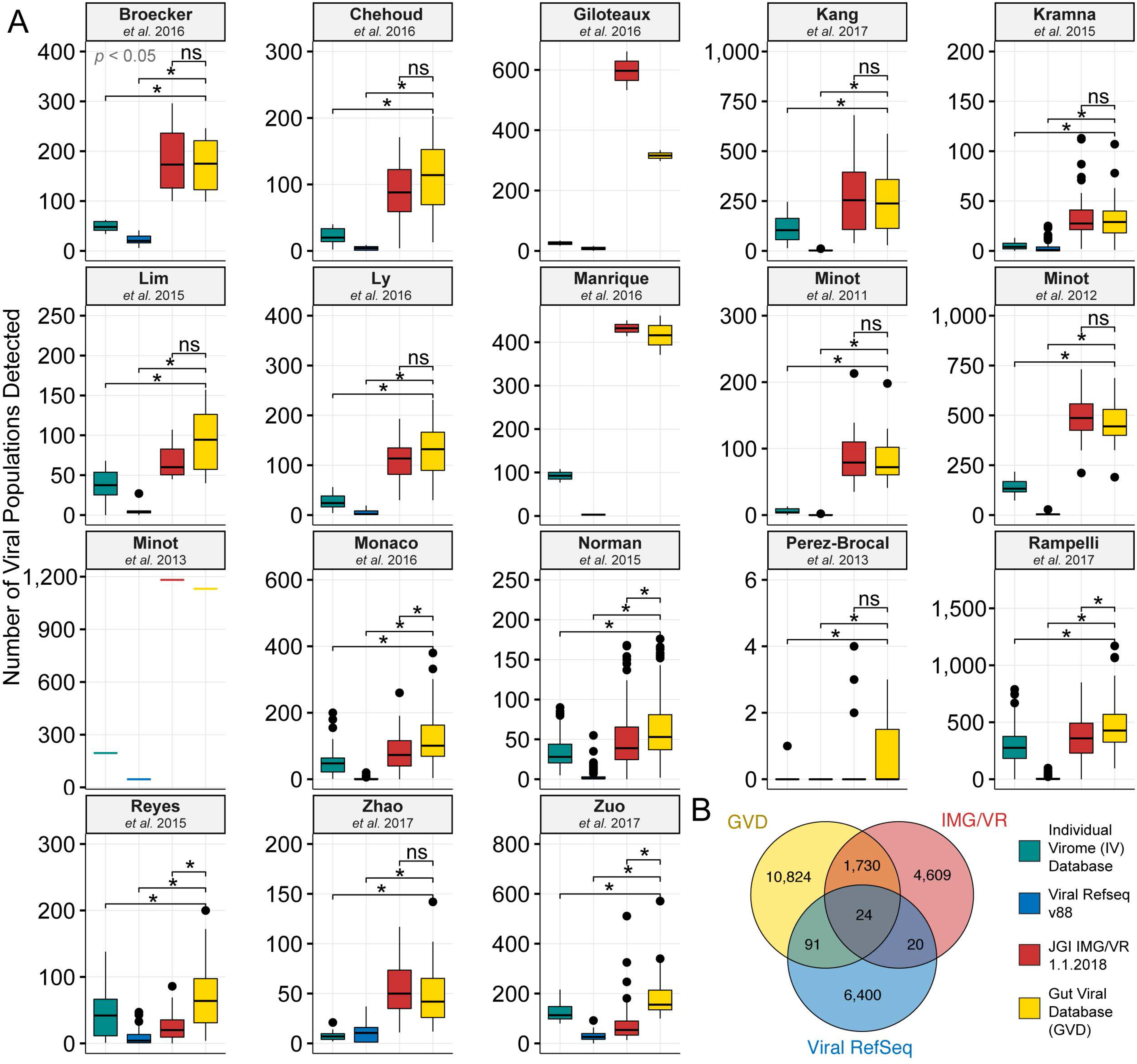
GVD as a reference database increases viral population detection. **(a)** Boxplots showing median and quartiles of the number of viral populations detected per study using the IV, Viral Refseq v88, JGI IMG/VR, or GVD databases. Studies where the reads were given to us prior to publication are excluded from this analysis (Yinda et al., 2019; Neto et al. (unpublished)). **(b)** Venn diagram showing the number of viral populations unique and shared between the different databases. Importantly, we only compared dereplicated viral populations from IMG/VR that came directly from human gut samples or had reads mapping to them from GVD gut samples.

### MDA amplification skews diversity and prohibits quantitative analysis of gut viromes

To evaluate this possible reduced viral “signal”, we first examined the role of methodological approaches in influencing inferences about ssDNA viruses. This is because we noticed that the bulk of ssDNA eukaryotic viruses (Anelloviruses, Circoviruses, Genomoviruses, Geminiviruses) and phages (Microviruses) originated from only 4 of the 21 studies gathered in this work (**Fig. 4 a,b**). These studies evaluated 2 infant gut viromes^19,37^ and 2 adult inflammatory bowel disease viromes^31,32^, and they reported relative abundance shifts of ssDNA and dsDNA phages within these viromes. From this observation, these studies concluded that such shifts could discriminate between healthy and disease states associated with virome development in early life.

**Figure 4.**
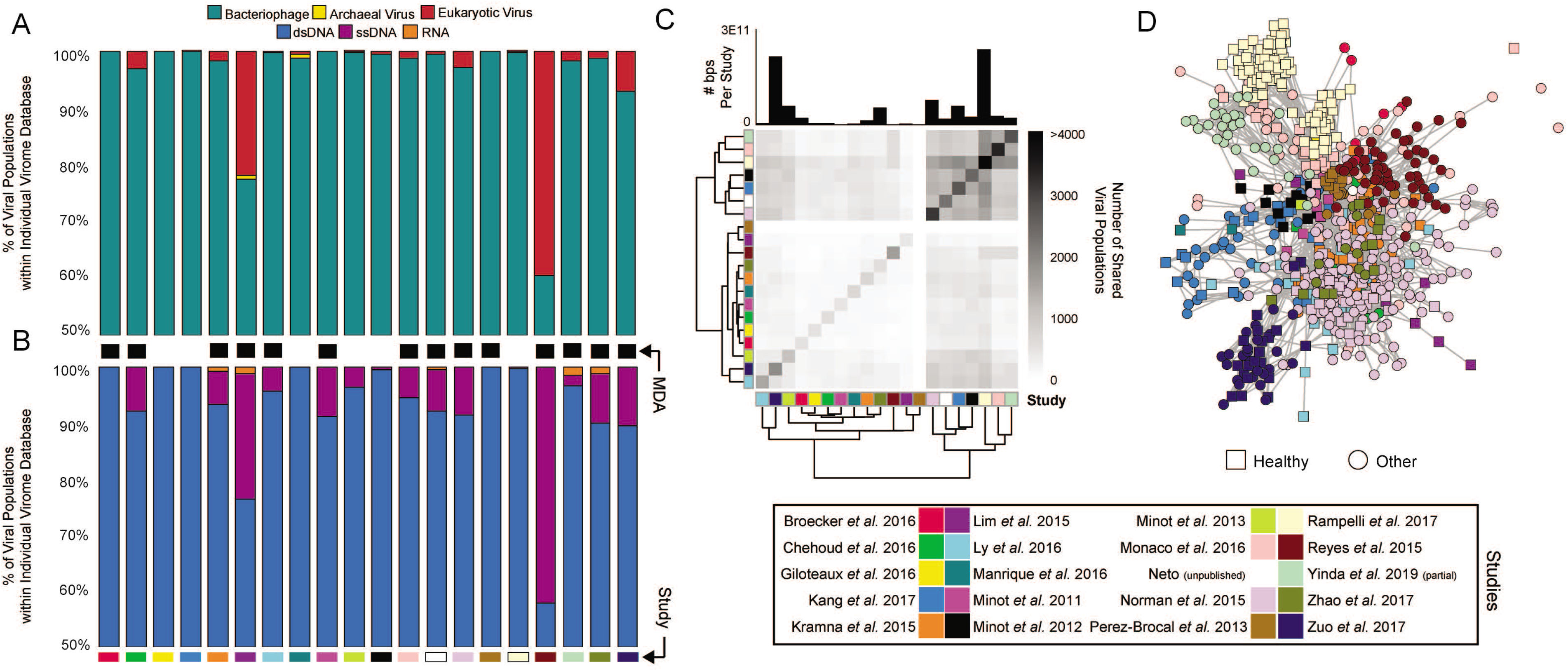
Individual Viromes (IV) Study Databases and Cross-Study Comparisons. **(a)** Barplot showing the proportion of those viruses that are bacteriophages, archaeal viruses, or eukaryotic viruses. The total number of assembled viral contigs and viral populations per study are available in **Supplementary Fig. 1a. (b)** Barplot showing the proportion of those viruses that are dsDNA, ssDNA, or RNA viruses. Studies where multiple displacement amplification (MDA) was used show a higher prevalence of ssDNA viruses. No viral contigs ≥1.5kb were assembled from the Reyes *et al*. 2010 study. **(c)** Hierarchically clustered heatmap showing the number of viral populations shared within and between studies. The barplot on top of the heatmap shows the total number of sequenced base pairs following quality control within each study. **(d)** Viral population co-occurrence network per individual within each study shows that individuals within a study cluster together regardless of health status. The squares represent the healthy individuals within each study.

However, the abundance of ssDNA viruses can also be enriched from methodologies used in making the viromes, even if all samples are processed consistently. Specifically, early virome studies where limiting viral nucleic acids were obtained, often used whole genome amplification kits that leverage a DNA polymerase from the phi29 ssDNA virus to obtain many-fold increases in DNA via multiple displacement amplification or MDA^54^. Though attractive at first, MDA is now known to have stochastic biases (e.g., 100s–10,000s-fold biases in coverage,^55,56^), which result from randomized initial template interactions and can induce chimera formation and uneven amplification of linear genomic sections (whether ssDNA or dsDNA templates), as well as systematic biases resulting from preferential amplification of small, circular and ssDNA genomes^57–61^. Taken together, MDA-associated artifacts skew the taxonomic representation of a community in non-repeatable ways and preclude quantitative analysis of viromes^57^. Although non-quantitative, MDA-amplified viromes do still have value enriching for ssDNA viruses, as well as estimating presence of viruses.

Consistent with the idea that these ssDNA viruses are methodologically enriched in the MDA libraries, we found that non-MDA amplified gut viromes contained significantly less ssDNA viruses than MDA amplified gut viromes (range: 0% - 4% versus 0-42%; Mann-Whitney U-test; *p*-value = 0.0083), though sample size was quite low. Further, while we see a strong linear relationship (R^2^ = 0.86) between sequencing depth and the number of viral populations sequenced in non-MDA viromes, this relationship is weak in MDA viromes (R^2^ = 0.39), suggesting that MDA can skew the number of assembled viral contigs in datasets (**Supplementary Fig. 1b**). Critically, 14 of the 21 studies gathered in this work employed MDA, which calls into question the quantitative nature of these datasets. Fortunately, viral nucleic acid extraction from feces often yield sufficient quantities for high throughput sequencing^26^, and in cases where they do not there are now several viable alternative methods to more quantitatively establish viromes with as little as 1pg of DNA^61,62^. Problematically, current established gut virome protocols recommend an MDA step^48,63^. If a researcher’s goal is to provide quantitative datasets, then we strongly advocate against this recommendation and instead suggest that alternative methods^61,62^ be used to generate gut viromes.

### Human gut virome study conclusions are more impacted by methodology than disease state

Given a systematically processed GVD, we next sought to determine whether global clustering patterns would emerge between study themes between all dataset used to build GVD. To this end, viral populations identified in this study were matched back to their respective datasets, and used in a co-occurrence network analysis (see **Methods**) to assess co-variation at two levels: between study datasets (**Fig. 4c**), and between viromes across all datasets (**Fig. 4d**). Between datasets, the fraction of shared viral populations was low (mean: 3% ±3%; **Fig. 4c**), except for 6 datasets that clustered together (hierarchical clustering bootstrap = 100%; **Fig. 4c**) and had a higher level of shared viral populations (>4-fold increase). Presumably, these elevated similarities across the 6 datasets may be due to deeper sequencing (Fig. 4d, top panel) that allowed deeper sequencing into the rare tail of viral populations among samples. A similar trend was observed when looking at the level of individuals within each study (**Fig. 4d**), where the co-occurrence network revealed close clustering between individuals derived from the same study, irrespective of geographical origin, health status and/or diet. This per study clustering implies that, taken together, these studies are not comparable likely due inconsistent sampling and extraction methodologies. We then investigated the prevalence of gut viral populations amongst all samples, so as to establish whether any viral populations were detected in all samples (i.e., a ‘core’ gut virome^22^). On average, 138± 170 (average ± SD; range: 0 to 849) viral populations were detected per sample, but not one viral population was found across all samples. We then explored deeper to detect whether subsets of the samples would reveal shared viral populations. We found that only 28 viral populations occurred in over 20% of the GVD samples. Most viral populations were detected in very few samples. In fact, >40% of the viral populations occurred in <0.5% of the samples and 98% of the viral populations occurred in <0.1% of the samples in GVD (**Fig. 5 a, b** and **Supplementary Table 3**). Further, we specifically looked at the prevalence of crAssphages, a well-recognized, multi-genera group of phages known to be widespread in gut viromes^64^ (**Fig.5 b, c**). While crAssphages are ubiquitous across the GVD samples, there was not one crAssphage viral population found universally, with the most widespread crAssphage population occurring in only 38% of samples. Importantly, when we looked at all healthy samples and healthy western samples specifically, still no shared viral populations were identified in all samples. (**Supplementary Fig. 2a, b**). Assuming samples were sufficiently sequenced, this may be indicative that individuals carry a unique ‘gut virome fingerprint’, even between twins, which is perhaps not surprising given recent suggestions of a similar ‘fingerprint’ for gut microbes (the ‘personal’ microbial microbiomes^65^). This apparent lack of core gut virome among individuals contrasts with a recent report^22^, in which overlapping patterns of phage genomes between 2 unrelated healthy individuals, as well as within a re-analyzed larger cohort^66^ revealed three levels of sharing patterns: (i) core (phage found in >50% of samples, (ii) common (phage found in >20-50% of samples), and (iii) unique (phage found in <20% of samples). Our analyses showed no viral populations shared above >50% of samples, thus bringing into question the presence of a ‘core’ virome as previously defined^22^, as well as a very limited ‘common’ virome (20-50% sharing across samples), in which we observed either 1% (all healthy; n=132) or 0.1% (all healthy Westerners; n=18) of GVD viral populations, similar to the 3% previously reported^22^ (see **Supplementary Table 4**). Likely, this discrepancy with our results could be attributed to how viruses were identified through read mapping. In the initial study reporting a core virome^22^, a virus was considered present if a single read mapped to a genome, a very permissive cut-off which does not take into account shared homologous regions between distinct viral populations. In this study, we considered a virus present if reads mapped 70% of the genome length (if genome is <5kb) or reads mapped at least 5kb of the genome (for genome >5kb in length) (see Methods). While our cut-off is more conservative, it better ensures that we are detecting the same viral population. Nonetheless, the idea of a core virome might still be an open question.

**Figure 5.**
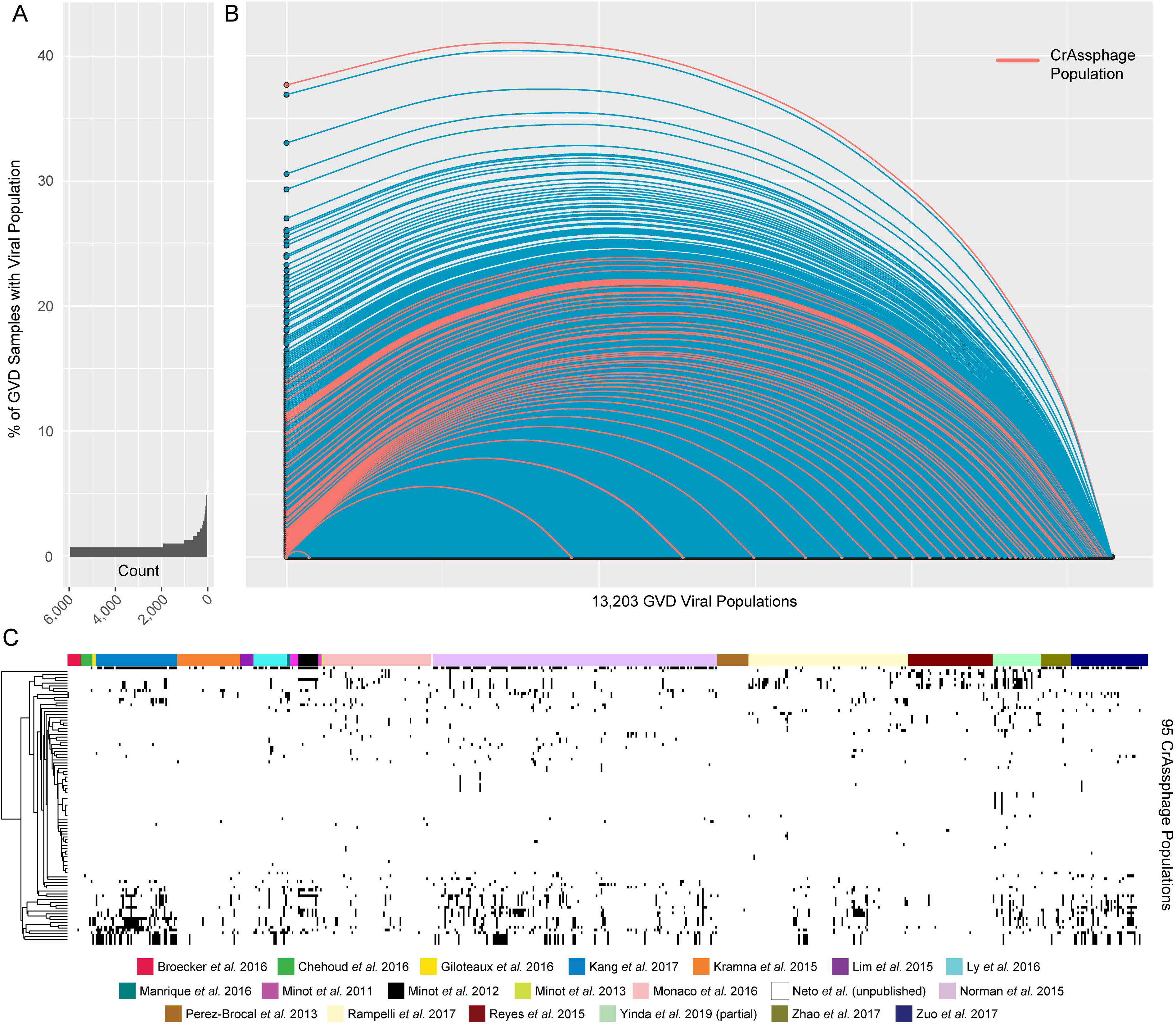
There are no core viral populations across GVD samples. **(a)** Histogram showing the number of viral populations present in different percentages of GVD samples. The vast majority of viral populations are found in <10% of the individuals. **(b)** Hive plot showing the percentage of GVD sample each viral population is detected within. The dots on the x-axis represent each GVD viral population in ascending order of the percentage of GVD samples that they are found within. The y-axis is the percentage of GVD samples that each viral population is detected within. CrAssphage viral populations are highlighted in red. **(c)** Heatmap showing the presence or absence of each crAssphage viral population across the different GVD samples.

### Re-evaluation of a previous study: the virome across different geographic regions and lifestyles

Due to the high level of sample clustering per study (**Fig. 4c**), we were unable to conduct cross-study analyses. Instead, we sought to assess if the virome community patterns between populations of varying lifestyles (industrialized versus semi-industrialized versus hunter-gatherer) would vary between the initial study^26^ or GVD-based, to test whether there were geographic biases around GVD viral populations, and how well sampled are the different geographic regions. This initial study encompassed a globally-distributed dataset (USA, Italy, Tanzania and two Peruvian populations: Tunapuco and Matses; **Fig. 6a**), and explored the impact of geography and diet on eukaryotic gut viruses (but did not include phages) and found that the hunter-gatherers (Hadza in Tanzania and the Matses in Peru) had the highest eukaryotic viral richness^26^.

**Figure 6.**
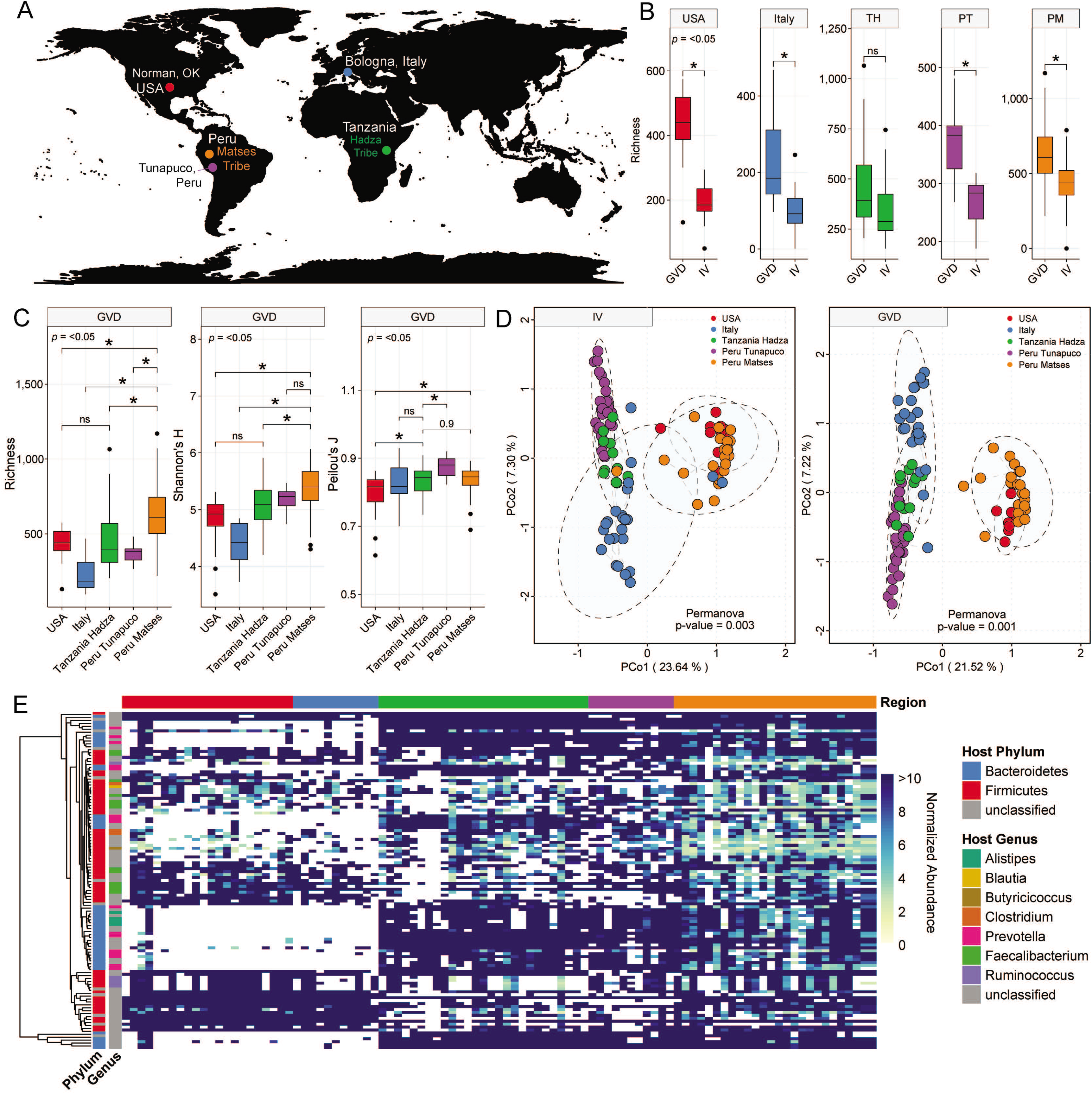
Diet and geography widely influence gut virome. **(a)** World map showing the geographical distribution of the Rampelli *et al*, 2017 dataset. **(b)** Boxplots showing median and quartiles of the number of viral populations detected using the GVD database and the Rampelli et al., 2017 viral database alone (IV) within each geographic group. **(c)** Boxplots showing median and quartiles of the α-diversity metrics – richness, Shannon’s H and Peilou’s J – across the different geographic groups using the GVD database (see **Fig. S3** for α-diversity metrics using the IV database). **(d)** Principal coordinate analysis (PCoA) of a Bray-Curtis dissimilarity matrix calculated from mapping the Rampelli et al., 2017 dataset against the IV (left) and GVD (right) databases. Analyses show that the viromes significantly (Permanova p < 0.05) structure into based on the geographic groups, with mapping to the GVD showing revealing much stronger clustering based on geography. Ellipses in the PCoA plot are drawn around the centroids of each group at a 95% confidence interval. The dashed lines connecting the different points reveal the connections determined by hierarchically clustering between the different samples. **(e)** Heatmap of the abundances of the viral populations found across >50% of individuals within the study. Individuals on a Western diet (from the USA and Italy) lack phages that infect Bacteroidetes, specifically those that infect *Prevotella* sp. All pairwise comparisons were performed using a two-tailed Mann-Whitney U-tests.

In this re-analysis, however, we included phages in addition to eukaryotic viruses, and focused on how the virome diversity varied along the dataset. We first evaluated whether per-region GVD-mediated detection of viruses would incur biases, potentially stemming from underrepresented viral populations from less-sampled geographical regions. This did not appear to be the case, as significant increases in virus detection were observed across 4 out of the 5 regions sampled (**Fig. 6b**). We next calculated diversity indices (**Fig. 6c** and **Supplementary Fig. 3**) for each regional dataset, and looked at the number of viral populations mapped with GVD. Overall, we reached a similar conclusion to the initial study (even when considering phages), in which the hunter-gatherers (Peru Matses) generally contained higher viral richness (**Fig. 6c - left**) and biodiversity (Shannon’s H, **Fig. 6c - middle**), but not higher evenness (Peilou’s J, **Fig. 6c -right**). Collector’s curves revealed that we have not saturated the human gut viral diversity among individuals globally (**Supplementary Fig. 4**) or even among just among American samples (**Supplementary Fig. 2, inset**). Thus, it appears much more viral diversity remains to be discovered across all geographic regions.

We next wondered whether the addition of phage in our analysis would reflect on overall viral community similarities by using Bray-Curtis distances between individuals across these geographic and lifestyle gradients (**Fig. 6d**). While unequal database representation can have an impact on alpha-diversity, beta-diversity is often less impacted^67^. Principal coordinate analyses (PCoA) of Bray-Curtis distances derived from using the individual Rampelli et al., 2017 virome database (**Fig. 6d**, left panel) and GVD (**Fig. 6d**, right panel) revealed no significant differences (Mantel’s test; R = 0.95, *p* = 0.001). However, analysis of the GVD-referenced PCoA revealed individuals with the same lifestyle and from the same region clustered together (PERMANOVA; *p* ≤ □0.001) and provided better resolution of the clustering in comparison to the IV-referenced PCoA. However, lifestyle alone may not account for the observed clustering patterns. The viromes of the Hadza in Tanzania and semi-industrialized, agrarian Tunapuco population in Peru strongly overlapped (hierarchical clustering bootstrap = 100%; **Fig. 6d**), most likely driven by their diets rich in root vegetables^68–70^. Nonetheless, when we look at differences between dominant viral populations (found in >50% individuals) across these geographic and lifestyle gradients, we see that there are key viruses missing from Western, industrialized gut viromes (**Fig. 6e**), specifically viruses that infect the genus *Prevotella spp*. This parallels the bacterial analyses that show that *Prevotella spp*. are enriched in non-Western gut microbiomes and many species are missing from Western, industrial gut microbiomes^69–71^. Overall, this suggests that lifestyle and diet has an impact not only on the bacterial community, but also on the viral community in the gut.

## CONCLUSIONS

The lack of a curated database for the detection of viral sequences in the human gut has been identified as the most critical shortcoming of applying metagenomic approaches to studying the human gut virome^72^. Although GVD is geared towards filling this gap and performs well (increasing viral detection 59-fold over the most commonly used database, NCBI viral RefSeq), there are limitations. *First*, the geographic and ethnic representation across the dataset is not very broad. Meta-analyses will benefit from more broadly representative datasets. *Second*, GVD was built using all datasets available by the end of 2017. Since then, as of May 2019, there are 11 additional datasets that study the gut virome, 8 of which use viral particle-enriched metagenomes (**Supplementary Table 5**). Further, there are many more human gut microbial metagenomic datasets and these could be a rich source for virus reference genomes as found for soils^73^ and the large-scale Earth Virome study^74^. To maintain significance as a resource, we will update GVD annually by extracting the viral signal from such gut-related datasets, as well as monitoring IMG/VR for gut-related viruses that should be integrated. *Third*, GVD is accessible through direct download as a single fasta file containing all GVD viral populations (see link in the ‘Data availability’ statement below), and is likely best paired with IMG/VR to maximize viral signal recovery. Future GVD updates and development will be required to improve the user experience for those not comfortable at command-line interfaces, but these are likely best integrated with large-scale standardizing efforts like the National Microbiome Initiative.

Given the relatively minimal value added via non-quantitative MDA-based approaches and the availability now of low-input quantitative approaches pioneered studying ocean viruses^61,75^ suggest that gut virome studies should move away from the former towards the latter. GVD, combined with the means to classify uncultivated virus genomes^47^, are prime starting requirements for enabling ecosystem-wide examinations^76^ of the dynamics and impacts of the virome within the human gut. Other environmental advances also invite such studies to include assessing the role of micro- and macro-diversity on virus persistance^41^, and metabolic reprogramming via virus-encoded auxiliary metabolic genes^73,76^. These combined efforts are critical to enable studies of the human gut virome to advance from ‘stamp collecting’ diversity studies towards the kinds of comprehensive efforts needed to incorporate viruses into mechanistic, predictive models. Such efforts, with future viral mapping outside the gut to parallel efforts for the ‘non-gut’ human microbiome^77^, should help transform personalized medicine and lead to a better understanding of human ecosystems.

---

**Supplementary Figure 1. Number of assembled viral contigs and populations. (a)** Barplot showing the number of assembled viral contigs versus the number of deduplicated viral populations per study. **(b)** Scatterplots with linear regressions showing the impact of increased sequencing on the number of assembled contigs per study divided by studies that did not have multiple displacement amplification (MDA; **top**) and those that did have MDA (**bottom**).

**Supplementary Figure 2. There are no core viral populations across healthy samples and across healthy western samples.** Hive plots showing the percentage of GVD samples each viral population is detected within across **(A)** all healthy individuals and **(B)** across only healthy western adults. The dots on the x-axis represent each GVD viral population in ascending order of the percentage of GVD samples that they are found within. The y-axis is the percentage of GVD samples that each viral population is detected within.

**Supplementary Figure 3.** Boxplots showing median and quartiles of the α-diversity metrics – richness, Shannon’s H and Peilou’s J – across the different geographic groups in the Rampelli *et al*. 2017 study using the IV and GVD databases.

**Supplementary Figure 4. The number of gut viral populations will still increase with more samples added to GVD.** Collector’s curve for gut viral populations in the GVD. (**inset**) Collector’s curve for just the viromes from samples from the USA.

**Supplementary Table 1.** Origin of datasets and associated metadata used to create the gut virome database.

**Supplementary Table 2.** Gut Viral Database contigs family-level taxonomy and putative hosts.

**Supplementary Table 3.** Distributions of viral populations across GVD samples.

**Supplementary Table 4.** Core, common, low-overlap, and unique GVD viral populations

**Supplementary Table 5.** Human gut virome studies since the end of 2017

## METHODS

### Experimental Model and Subject Details

Gut virome database (GVD) studies were selected by doing a thorough and manually curated search of the Web of Science Core Collection of Thomson Reuters for studies looking at viruses in the gut published prior to 2018. All studies that used next-generation sequencing and looked for viruses within the gut microbiome were selected to be part of GVD (see full list of studies in **Supplementary Table 1**). Additionally, we were given access to the reads of two studies that were unpublished at the time. One of the studies, however, is now published (Yinda et al., 2019).

### Viral contig assembly, identification, and dereplication

Previously published GVD reads were downloaded from their respective hosting databases (e.g. SRA, iVirus, or MG-RAST). Prior work revealed that an individual’s gut virome is stable across time (Minot et al., 2013), so reads were pooled per individual regardless of the number of time points, with a few exceptions (**Fig. 1**). These exceptions included studies with fecal microbiota transfers and studies whose reads were given to us prior to publication. For fecal microbiota transfers, all time points per individual were kept separate and processed independently. Read sets from two studies were given to us prior to publication (Yinda et al., 2019; Neto et al., unpublished). For the Yinda *et al*., 2019 study, individual’s reads were pooled based on their location, age, and contact with bats. The pools were counted as a single individual’s virome for our analyses. For the Nadia et al., (unpublished), all reads from all individuals were pooled together. A global map showing the number of individuals (or pooled read sets) originating from each country was created using the R packages ‘rworldmap.’ In total, there were 648 GVD samples from 572 individuals.

Pooled reads were then assembled using metaSPAdes 3.11.1^78^. Following assembly, contigs ≥1.5kb were piped through VirSorter^79^ and VirFinder^80^ and those that mapped to the human, cat or dog genomes were removed. For viral-enriched metagenomes (i.e. viromes), contigs ≥5kb or ≥1.5kb and circular that were sorted as VirSorter categories 1-6 and/or VirFinder score ≥0.7 and *p* <0.05 were pulled for further investigation. Of these contigs, those sorted as VirSorter categories 1 and 2, VirFinder score ≥0.9 and *p* <0.05 or were identified as viral by both VirSorter (categories 1-6) and VirFinder (score ≥0.7 and *p* <0.05) were classified as viral. The remaining contigs were run through CAT^81^ and those with <40% (based on an average gene size of 1000) of the genome classified as bacterial, archaeal, or eukaryotic were considered viral. For the microbial metagenomes, we took a more conservative approach with only contigs ≥5kb or ≥1.5kb and circular that were sorted as VirSorter categories 1-2 and VirFinder score ≥0.6 and *p* <0.05 were considered viral. Across the both the viral-enriched and microbial metagenomes, contigs ≥5kb or ≥1.5kb and circular that were classified as eukaryotic viral contigs by CAT were also considered viral. In total, 29,345 viral contigs were identified.

Viral contigs that were from known ssDNA or RNA viral families using CAT were grouped into populations if they shared ≥95% nucleotide identity across ≥100% of the genome. Because there are no benchmarked metagenomic population boundaries for ssDNA and RNA viral families, we chose to not use stringent dereplication. All other contigs were considered double-stranded DNA and were grouped into populations if they shared ≥95% nucleotide identity across ≥70% of the genome (*sensu*^82^) using nucmer^83^. All the viral contigs that were assembled were dereplicated per study to create the individual virome (IV) databases and across all of GVD (see **Supplementary Fig. 1**). For GVD, this resulted in 13,203 total viral populations found in GVD (see **Supplementary Table 3** for VirSorter, VirFinder, and CAT results), of which 6,373 were ≥10kb in length.

### Core Viral Population Analyses

To explore if there were any core viral populations, the abundance table was turned into a binary presence-absence matrix. The number of GVD samples that each viral population was detected within was then calculated using R and divided by the total number (648) to get the percentage of samples. Each viral population’s percentage was plotted in hive plot using ‘geom_curve’ in ggplot2^84^. This process was repeated on subsets of the matrix including all healthy individuals and only the healthy western adults. The number of viral populations that were present across different percentages were calculated using R and their distributions plotted using ‘geom_histogram’ in ggplot2^84^. CrAssphage viral populations in GVD were identified using CAT results and by dereplicating GVD viral populations with the crAssphage genomes identified in Guerin *et al*^6^ and seeing which GVD genomes cluster. In total, there were 95 unique crAssphage populations. The binary presence-absence data for the crAssphage populations were plotted using pheatmap in R.

### Viral taxonomy

For each viral population, ORFs were called using Prodigal^85^ and the resulting protein sequences were used as input for vConTACT2^47^ and for BLASTp. Double-stranded DNA viral populations represented by contigs >10kb were clustered with Viral RefSeq release 88 viral genomes using vConTACT2. Those that clustered with a virus from RefSeq based on amino acid homology based on DIAMOND^86^ alignments were able to be assigned to a known viral taxonomic genera. For viral dsDNA populations that could not be assigned taxonomy or were <10kb, family level taxonomy was assigned using a majority-rules approach, where if >50% of a genome’s proteins were assigned to the same viral family using a blastp bitscore ≥50 with a Viral RefSeq virus, it was considered part of that viral family (see **Supplementary Table 3** for family-level taxonomy). For ssDNA and RNA viruses, CAT was used to assign the viral family (see **Supplementary Table 3** for family-level taxonomy).

### Viral Host Prediction

Bacteriophage hosts were predicted using a variety of bioinformatic methods including: (i) CRISPR-spacer matches, (ii) prophage blasts, (iii) tRNA genes matches, and (iv) WiSH matches^87^ against Bacterial Refseq v88. CRISPR spacers were predicted using MinCED (https://github.com/ctSkennerton/minced) and the CRISPR Recognition Tool (CRT^88^) and a BLASTn (-task blastn-short -word_size 5) was used to assess matches between the CRISPR spacers and viral populations in GVD. Those with 1 mismatch were considered a match. For prophage blasts, a blastn of the viral population against Bacterial RefSeq was performed. A bacterial genome with ≥2500bp regions of their genome matching at 95%ID with a viral population genome were considered putative hosts of that viral population (see^76^). Viral tRNA genes and Bacterial RefSeq tRNA genes were predicted using tRNA-scan^89^ and then a blastn was performed between the viral and bacterial tRNA genes. Bacterial tRNA genes that matched viral tRNA genes at 95% ID across 100% of the length were considered putative bacterial hosts. Lastly, WIsH was used to predict hosts according to default settings^87^. Priority host assignment was given to CRISPR, then prophage, WIsH and tRNA results. Viruses with putative archaeal hosts were predicted using MarVD^90^. Viruses with predicted eukaryotic hosts were assigned based on their assigned taxonomic viral family.

### Detecting viral populations and calculating their raw abundances

To calculate the raw abundances of the different viral populations in each sample, reads from each GVD pooled read set were first non-deterministically mapped to all GVD viral population genomes using bowtie2. Further, reads from each GVD pooled read set per study were mapped to their respective IV databases. BamM (https://github.com/ecogenomics/BamM) was used to remove reads that mapped at <95% nucleotide identity to the contigs, bedtools genomecov^91^ was used to determine how many positions across each genome were covered by reads, and custom Perl scripts were used to further filter out contigs without enough coverage across the length of the contig. All contigs ≤5kb in length with >70% of the contig covered were considered detected in the sample. Contigs >5kb in length with ≥5kb in length covered were also considered detected in the sample^92^. BamM was used to calculate the average read depth (‘tpmean’ - minus the top and bottom 10% depths) across each detected contig. For the alpha-diversity calculations, the average read depth was used as a proxy for abundance and normalized by total read number per metagenome to allow for sample-to-sample comparison. However, because most of the studies in GVD involved MDA, which can skew abundances, we chose to use only a presence-absence statistic (richness) for most of our α-diversity calculations. Collector’s curves and the whole GVD and across only American samples were calculated using the function ‘specaccum’ in the R ‘vegan’ package^93^.

### Comparisons to IMG/VR, Viral RefSeq v88, and IV databases

The IMG/VR (1.1.2018 release) included all viral contigs assembled from different datasets. All of the viral contigs in GVD, Viral Refseq v88, and IV databases are dereplicated at the population level. In order to make IMG/VR comparable to GVD, Viral Refseq and IV databases, we needed to dereplicate the IMG/VR database. IMG/VR (1.1.2018 release) is composed of 715,672 contigs. Because dereplication is extremely computationally intensive, we decided to only focus on dereplicating viral contigs that originated from the human gut and had at least 1 read from a GVD metagenome map. These IMG/VR viral contigs were then dereplicated using the same methodology as previously described in the methods section. In total, 29,378 IMG/VR viral contigs were dereplicated into 6,652 viral populations. GVD pooled read sets were mapped to this IMG/VR human gut viral population database, Viral RefSeq v88, and the IV databases for each individual study in GVD. The raw abundances of the different IMG/VR and Viral RefSeq viral populations in each sample were calculated the same way as described in the previous section. The total number of viral populations detected per sample per study using the different databases were then plotted and comparative statistics using the ‘ggboxplot’ function from the ‘ggpubr’ package in R.

All of the viral populations from GVD, the dereplicated IMG/VR gut-specific dataset, and Viral Refseq were then dereplicated to see how many viral populations overlapped between databases. The results were then plotted using the ‘VennDiagram’ package in R. Importantly, in the dereplication process, some of the original viral populations in each database may be dereplicated down due to the presence of a longer viral contig from the same population that links the two together into the same population. Across the databases, 329, 177, and 459 viral populations were dereplicated in GVD, IMG/VR, and Viral Refseq, respectively. This is why the total number of populations displayed in the Venn diagram does not add up to the total number of viral populations in each database.

### Clustering studies based on shared viral populations

To test how studies clustered together, the viral population presence-absence data from individuals (or pooled read sets) within a study were merged. In Study 1, individual A had viral population 1, 2, 4, 5 and individual B had viral population 3, then Study 1 had viral populations 1, 2, 3, 4, and 5. The different studies were then assessed for the number of shared viral populations that were present in both studies. These values were then displayed and hierarchically clustered using the R ‘pheatmap’ package and the stability of the hierarchical clusters were assess using the R ‘pvclust’ package. The number of shared viral populations between individuals (or pooled read sets within a sample) were clustered using the R ‘SPIEC-EASI’ package^94^ using the Meinshausen and Buhlmann (MB) method to infer associations between samples based on the shared number of viral populations. The network was plotted using the R ‘igraph’ package.

### Alpha- and Beta-Diversity calculations

The α- (Richness, Shannon’s *H*, and Peilous’ *J*) and β- (Bray-Curtis dissimilarity) diversity statistics were performed using VEGAN^93^ in R. For all studies, except for Rampelli et al.^26^, only richness was calculated for both abundances based on read mapping to IMG/VR, Viral Refseq, the IV databases and GVD. Comparisons were plotted using ‘ggboxplot’ function in the R ‘ggpubr’ package. The Rampelli et al.^26^ did not use MDA, so we went ahead with scaling the raw abundances based on the number of quality controlled base pairs sequenced to normalize the data. All α-diversity statistics were calculated and β-diversity was used to look at community structure using both the IV and GVD databases. Principal Coordinate analysis (function capscale of VEGAN package with no constraints applied) was used as the ordination method to plot the Bray-Curtis dissimilarity matrices (function vegdist; method “bray”) after a cube root transformation (function nthroot; n = 3). To determine if the Rampelli et al. samples clustered by geographic region, a permanova test (function “adonis’’) and the 95% confidence interval were plotted using function “ordiellipse.” Further, the samples were hierarchically clustered and plotted within the PCoA. To specifically look at abundance differences in the most abundant viral populations in the Rampelli et al.^26^ study, viral populations that were present in 50% study individuals and their hosts information were plotted using the R ‘pheatmap’ package.

## Code availability

Scripts used in this manuscript are available on the Sullivan laboratory bitbucket under Gut_Virome_Database.

## Data availability

All raw reads are available through SRA, iVirus, or MG-RAST using the identifiers listed in **Supplementary Table 1.** GVD viral populations can be downloaded directly from iVirus through the following link: https://de.cyverse.org/dl/d/E83EFBFF-2A23-4794-8819-ADD34160D018/FINAL_Gut_Viral_Database_GVD_1.7.2018.fna

## ACKNOWLEDGEMENTS

Computational support was provided by an award from the Ohio Supercomputer Center (OSC) to MBS. Study design and manuscript comments from Shini Sunagawa, Miguelangel Cuenca Vera, Bas E. Dutilh, Ksenia Arkhipova, Pedro Meirelles and Simon Roux are gratefully acknowledged. Funding was provided by the Gordon and Betty Moore Foundation (#3790) to MBS and an NIH T32 training grant fellowship (AI112542) to ACG.

## AUTHOR CONTRIBUTIONS

A.C.G. collected all datasets and metadata for the study. A.C.G. and A.H. curated metadata for the study. A.C.G., O.Z., A.H., B.B., M.B.S created the study design, analyzed the data, and wrote the manuscript. All authors approved the final manuscript.

## MATERIALS & CORRESPONDENCE

Correspondence and material requests should be addressed to Matthew B. Sullivan at.

