## Supplementary figures and images for "The human gut virome database"

### Supplemental Data 1

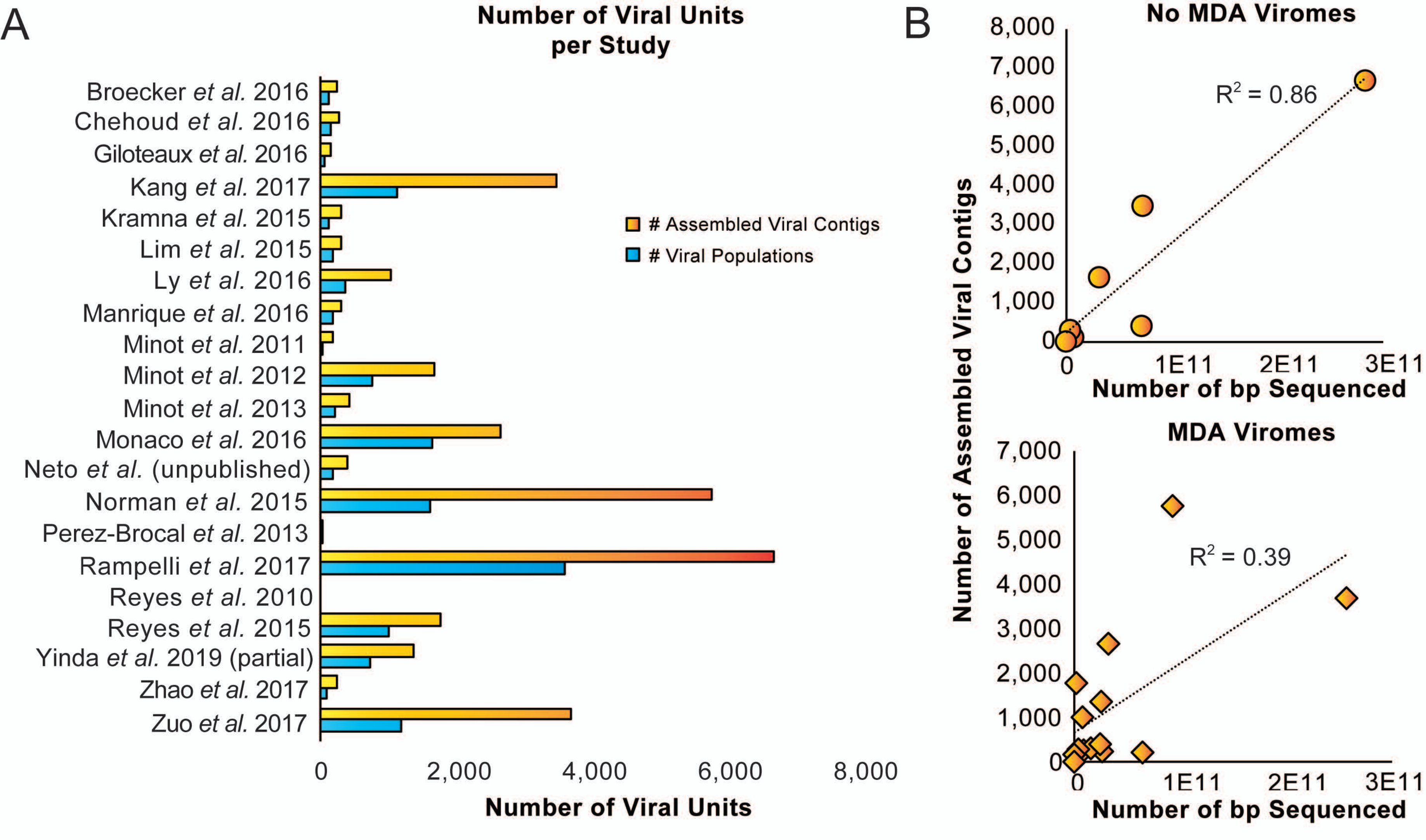

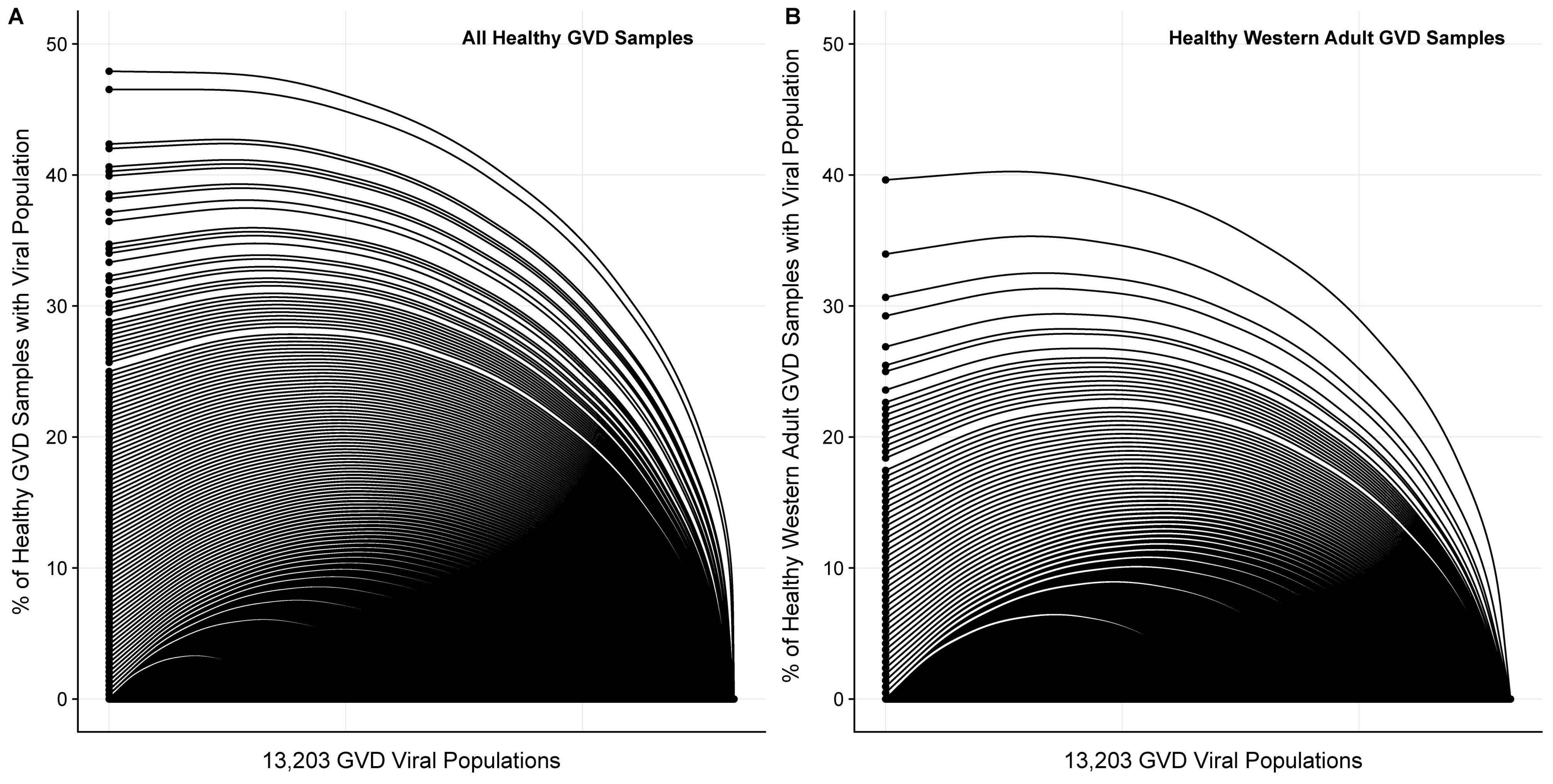

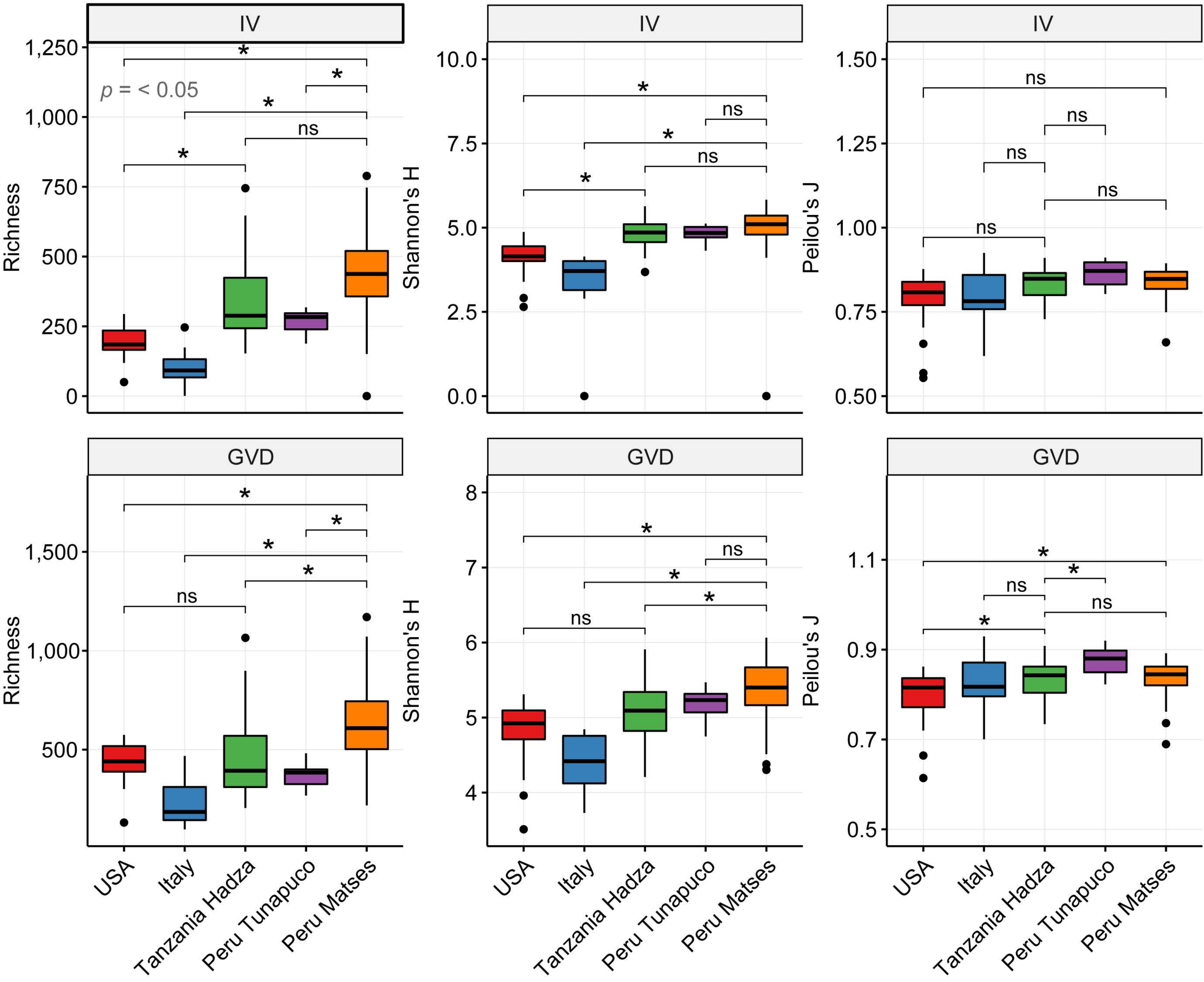

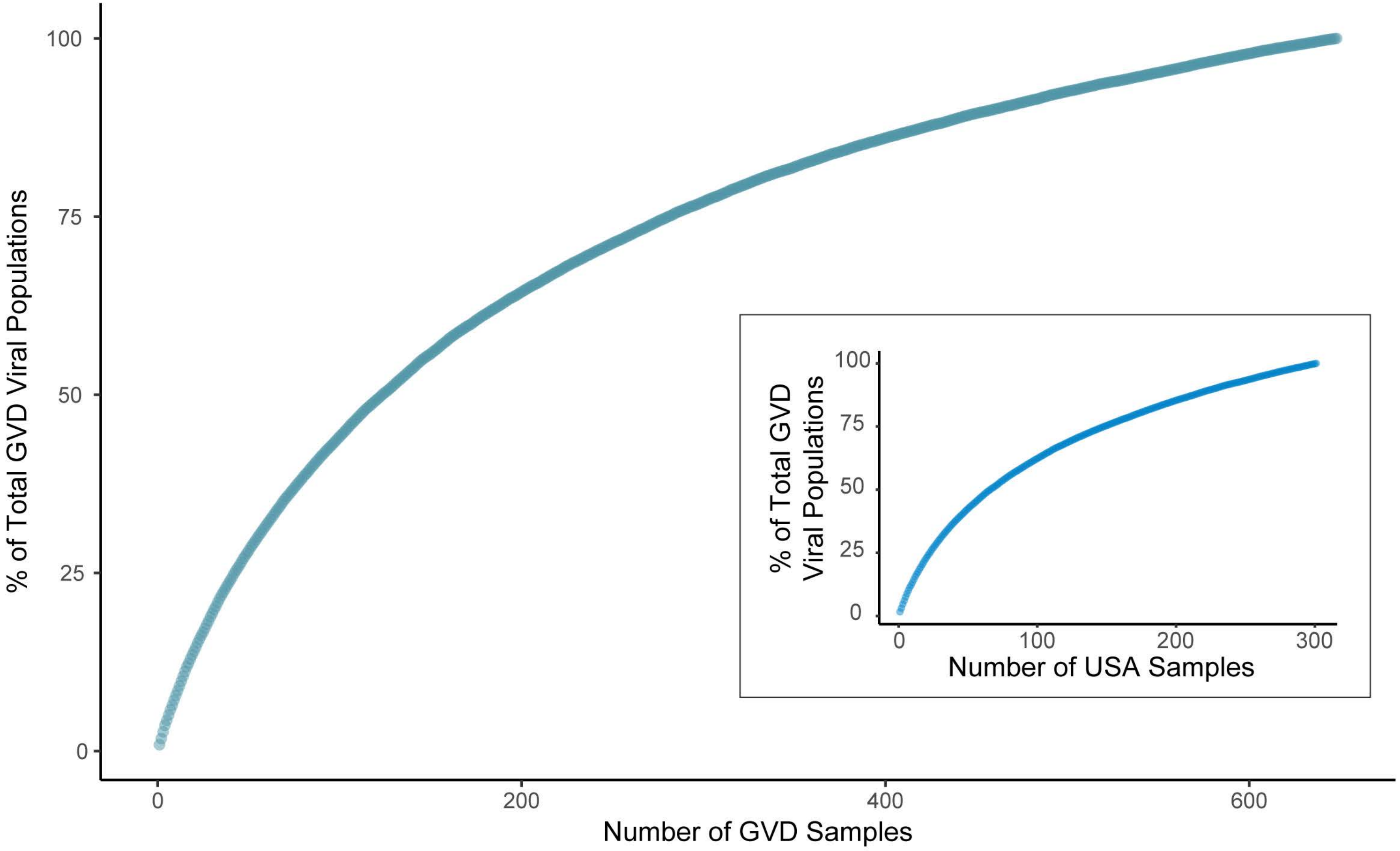
